# CERI, CEFX, and CPI: largely improved positive controls for testing antigen-specific T cell function in PBMC compared to CEF

**DOI:** 10.1101/2020.11.12.380188

**Authors:** Alexander A. Lehmann, Pedro A. Reche, Ting Zhang, Maneewan Suwansaard, Paul. V. Lehmann

**Affiliations:** Cellular Technology Ltd., Shaker Heights, OH, United States; Laboratorio de Inmunomedicina & Inmunoinformatica, Departamento de Immunologia & O2, Facultad de Medicina, Universidad Complutense de Madrid, Madrid, Spain

**Keywords:** T cell immune monitoring, CD8+ T cell immunity, CD4+ T cell immunity, antigen presenting cell functionality, PBMC fitness, immune dominance, T cell determinant, aleatory T cell recognition, ELISPOT, FluoroSpot, ImmunoSpot, SARS-CoV-2, COVID-19

## Abstract

Monitoring antigen-specific T cell immunity relies on functional tests that require T cells and antigen presenting cells to be uncompromised. Drawing of blood, its storage and shipment from the clinical site to the test laboratory, and the subsequent isolation, cryopreservation and thawing of peripheral blood mononuclear cells (PBMC) before the actual test is performed can introduce numerous variables that may jeopardize the results. Therefore, no T cell test is valid without assessing the functional fitness of the PBMC being utilized. This can only be accomplished through inclusion of positive controls that actually evaluate the performance of the antigen-specific T cell and antigen presenting cell (APC) compartments. CEF peptides have been commonly used to this extent. Here we show that CEF peptides fail as a positive control in nearly half of test subjects. Moreover, CEF peptides only measure CD8+ T cell functionality. More reliable alternatives for the assessment of CD8+ T cells are introduced here, as well as positive controls for the CD4+ T cell and APC compartments. In sum, we offer new tools and strategies for the assessment of PBMC functional fitness required for reliable T cell immune monitoring.

## 1. Introduction

For decades, immune diagnostics primarily focused on measuring antigen-specific serum antibody reactivity. Until recently, measuring antigen-specific T cells has been limited to animal models and exploratory human studies. Presently, there is a surge of interest in T cell immune monitoring for humans, especially since the presence of antigen-specific antibodies is not a predictor of T cell-mediated effector functions, while the latter can be decisive for clinical success. Such is the case, for example, with therapeutic unleashing of T cells against tumors [1], or with the immune system’s ability to control the SARS-CoV-2 virus [2].

Several obstacles needed, and in part still need, to be overcome for reliable measurement of antigen-specific T cell immunity in humans. First, T cell monitoring relies on functional test systems. Antibodies in isolated serum are stable for years and this fact largely facilitates monitoring of humoral immunity. In contrast, while long-lived *in vivo* [3], T cells in blood are perishable and start dying shortly after their isolation from the body [4]. In most cases, the blood drawn at clinical sites needs to first be transported to a test laboratory where the PBMC containing the T cells and APC essential for functional T cell assays are isolated. Excessive shear forces exerted during drawing of the blood, a delay in its transportation, and exposure to extreme temperatures during transit can each cause damage to the T cells and APC, leading to an impairment of their fitness when tested [5]. As it is not practical to test the PBMC samples one by one as they arrive in the laboratory, most choose to cryopreserve these cells to enable later testing in larger batches, and/or to be able to repeat test results or to extend testing as needed. During freeze-thawing, T cells and APC can also incur damage. Indeed, the development of protocols to freeze and subsequently thaw PBMC without impairing antigen-specific CD4+ or CD8+ T cell functionality was one of the major milestones that enabled T cell immune monitoring [6].

With all the possible sources of damage to PBMC prior to performing the actual test, T cell assays are inconceivable without proper controls to verify the functional fitness of these cells. Establishing the ratio of live/dead/apoptotic cells in the PBMC prior to testing them is helpful yet insufficient to identify their fitness [7]. The functionality of antigen-specific T cells can only be established by measuring exactly that, which in turn requires positive control antigens to which ideally all humans can be expected to have developed T cell immunity. This article is dedicated to the study of such positive control antigens.

The second general obstacle to T cell immune monitoring is that antigen-specific T cells occur in low, and frequently very low, frequencies amongst all T cells in PBMC. This also applies to viruses to which humans have developed immunity, and which therefore could be suited as positive controls [8]. In subjects naïve to an antigen, the frequencies of antigen-specific T cells are typically less than 10 in 10^6^ within PBMC [9]. Moreover, although naïve T cells can recognize antigens, they do not have effector functions. Consequently, naïve T cells are typically undetectable in functional assays when tested directly *ex vivo*. During the course of an immune response, antigen-specific T cells (and B cells) clonally expand, however, and their frequency rises in the body and within PBMC. The numbers peak between one and two weeks after an infection/vaccination, at which time antigen-specific T cells can constitute up to 1% of the PBMC; subsequently, the frequencies rapidly decline before reaching a steady state in the range of 20-100 antigen-specific T cells per 300,000 PBMC (corresponding to 0.007-0.035%) [10]. Among the techniques presently available for reliably detecting such rare antigen-specific T cells, ELISPOT assays have become the forerunner. Subsequently, most efforts investigating the immune response elicited by candidate SARS-CoV-2 vaccines have relied upon ELISPOT to monitor the induced T cell response [11]. Likewise, this study also relies upon ELISPOT analysis for detecting antigen-specific T cells within PBMC.

In ELISPOT and the related FluoroSpot assays, antigen-specific T cells are visualized by detecting the cytokines they release following exposure to the test antigen: these cytokines are captured around each secreting cell on a membrane that has been pre-coated with cytokine-specific antibodies [12]. Thus, the secretory footprint of each antigen-specific T cell is retained on the membrane in the form of a cytokine “spot”. The subsequent visualization of these plate-bound cytokine “spots” permits one to count the number of test antigen-specific T cells (expressed as “spot forming units” or SFU) present within all PBMC plated in a well. In this way, the frequency of antigen-specific T cells, and thus the magnitude of antigen-specific T cell immunity, can be established [13]. Measurements of multiple cytokines simultaneously, either in double-color ELISPOT or multi-color FluoroSpot assays, can also define the effector lineage(s) of the antigen-specific T cells [14]. As is the case for any functional T cell assay, ELISPOT assays are also critically dependent on the functionality of the T cells and APC being preserved after storage/shipment of the blood, isolation of PBMC, and freezing and thawing of the cells before the actual test is performed [15].

The third obstacle to T cell immune monitoring is that the choice of the antigen/peptide that is used in any functional T cell assay will define whether the memory T cells that have been induced *in vivo* will be detected at all. For immune monitoring with exogenous protein antigens this is not an issue, but such will largely detect only CD4+ T cells, and not CD8+ T cells [16]. When exogenous protein antigens are added to PBMC, professional APC (macrophages, dendritic cells and B cells) will process and present the antigen. The APC takes up the protein antigen, degrades it, and loads it onto MHC class II molecules (but not efficiently onto MHC class I molecules). The APC then transports the peptide-loaded class II molecule to its cell surface for antigen presentation to CD4+ T cells. When protein antigens are used for T cell recall assays, it is therefore not important to know the HLA class II alleles expressed by the test subject, nor the exact peptide epitope that is being presented to the CD4+ T cells: natural antigen processing and presentation mechanisms inherent to the APC select the relevant epitopes for each test subject without involving human estimation. Equally importantly, when using protein antigens, no epitope will be left behind, but the full antigen-specific CD4+ T cell repertoire induced *in vivo* during the immune response will be detected *in vitro* in the recall assay.

Reliably detecting antigen-specific CD8+ T cells in PBMC is much more intricate. CD8+ T cells evolved to survey ongoing protein synthesis within cells of the body, thus permitting CD8+ T cells to identify virally-infected or malignant cells, so as to kill them. During protein synthesis within every cell, defective byproducts arise and are quickly degraded by the proteasome into peptide fragments. Some of these peptides are transported to the endoplasmic reticulum where they are loaded onto nascent HLA class I molecules and such peptide-loaded class I molecules are transported to the cell surface where they are displayed for recognition by CD8+ T cells [17]. As the HLA gene complex is polygenic and highly polymorphic, and each allelic HLA class I molecule has a unique peptide binding specificity [18], this natural antigen presentation process results in a unique array of peptides presented in each individual, that is dictated by the unique HLA allele composition in said individual. To further complicate things, not all peptides presented will elicit a CD8+ T cell response *in vivo:* even HLA allele-matched individuals – that should present the same peptide epitopes when infected with the same virus - frequently develop aleatory CD8+ T cell response patterns *in vivo* for the individual epitopes [19]. Exogenously added protein antigens are also inefficiently presented to CD8+ T cells in the context of class I antigens and require that the antigen be added to the test PBMC in the form of short peptides, 8-11 amino acids long, that can directly bind to HLA class I molecules. However, the usage of such peptides requires (a) customization of the peptides to the HLA class I alleles expressed by each test subject, and (b) a precise knowledge of which epitope will be presented by which class I allele. Major progress has been made regarding allele-specific epitope prediction[20], but we are still far from customizing peptides for the reliable assessment of CD8+ T cell immunity [19,21,22]. However, selecting a single predicted peptide for immune monitoring that is not actually targeted by CD8+ T cells in an individual will result in false negative results[23]. Therefore, immune monitoring of CD8+ T cells increasingly moves towards usage of large peptide libraries in an attempt to cover all possible CD8+ T cell epitopes, rather than relying on individualized predictions of peptides [24].

Presently the CEF peptide pool is the gold standard to test the functionality of antigen-specific T cells, and as such is typically included as the positive control in most T cell assays: it originally consisted 23 well-defined CD8+ T cell epitopes of CMV, EBV, and Flu virus that have been selected to match HLA class I alleles that are frequent in Caucasians[8]. It was soon realized that these 23 peptides were insufficient to recall CD8+ T cells in most subjects, and so the number of peptides was increased to 32; constituting the extended CEF pool [25]. There remains an urgent need to further improve this positive control, as even the extended CEF pool frequently fails with PBMC that yield strong recall responses to CMV, EBV, and Flu virus [26]. In such cases, clearly, the negative CEF result does not signify impaired fitness of the PBMC, but suggests that CEF itself is not a reliable positive control. We tested here the hypothesis that the reason for CEF’s frequent failure to serve as a positive control is that 32 peptides of three viruses are not sufficient to reliably recall memory T cells in most human subjects, and that substantially broader epitope coverage is required. This hypothesis was directly tested by identifying CEF-low and non-responder subjects and evaluating their responses to extended pools representing common viruses that circulate in the human population.

## 2. Materials and Methods

### 2.1

PBMC from 210 randomly selected healthy human donors were obtained from the ePBMC library of Cellular Technology Limited (CTL, Shaker Hts, OH, USA, Cat# *CTL*-IP1). The HLA class I type, age, sex, and race for these donors is specified in Supplemental Table 1. These donors were recruited by Hemacare (Van Nuys, CA, USA) and the PBMC were isolated by leukapheresis at Hemacare using Hemacare’s IRBs. The PBMC were cryopreserved following protocols that maintain full T cell and APC functionality upon thawing [6], and were stored in liquid nitrogen vapor until testing. Thawing, washing, and counting of the cryopreserved cells was done as previously described [7]. Within 2 h after thawing, the cells were transferred into the ImmunoSpot^®^ assay.

### 2.2. CD4 and CD8 Depletion of PBMC

CD4+ and CD8+ T cell subsets were depleted from PBMC using magnetic bead-based CD4 and CD8 negative selection kits (Stem Cell Technologies, Vancouver, Canada). The cell separations were performed according to the manufacturer’s instructions.

### 2.3. Positive Control Antigens

#### 2.3.1.

CEF, a pool of 32 well-defined HLA class I-restricted epitopes of <u>C</u>MV, <u>E</u>BV, and <u>F</u>lu virus as defined in [25]. These peptides are 8-11 amino acid long and have been selected to recall CD8+ T cells. They were from, and are available through CTL (Catalog # CTL-CEF-002).

#### 2.3.2.

CEFX, by JPT Peptide Technologies, Berlin, Germany (Product Code: PM-CEFX) is a pool of 176 known peptide epitopes for a broad range of HLA sub-types – class I and class II – and different infectious agents, namely *Clostridium tetani*, Coxsackievirus B4, *Haemophilus influenza, Helicobacter pylori*, Human adenovirus 5, Human herpesvirus 1, Human herpesvirus 2, Human herpesvirus 3, Human herpesvirus 4, Human herpesvirus 5, Human herpesvirus 6, Human papillomavirus, JC polyomavirus, Measles virus, Rubella virus, *Toxoplasma gondii*, and Vaccinia virus. These peptides are 9-15 amino acids long, and have been selected to recall both CD4+ and CD8+ T cells. CEFX was tested at 1 μg/mL.

#### 2.3.3.

CPI: protein antigens of <u>C</u>MV, <u>P</u>arainfluenza and Influenza viruses, as described in [26]. CPI was from, and is available through CTL, Catalog #CTL-CPI-001. CPI was tested at 6.25 μg/mL.

#### 2.3.4.

CERI: 124 peptides of <u>C</u>MV, <u>E</u>BV, <u>R</u>SV, and Influenza virus. The individual peptides, 9 amino acids long, were selected based on peptide binding predictions for a broad range of HLA class I alleles expressed in all human races, and diverse ethnic subpopulations [27]. CERI was from, and is available through CTL, Catalog # CTL-CERI-300. CERI was tested at 1 μg/mL.

#### 2.3.5.

Anti-CD3 antibody was from Sigma-Aldrich, St. Louis, MO (clone OKT3, catalog # SAB4700040-100UG). It was tested at 0.05 μg/mL.

### 2.4. Human Interferon-γ ImmunoSpot^®^ Assay

The human interferon-γ (IFN-γ) ImmunoSpot^®^ test kits were used from CTL (catalog# hIFNgp-1M/10), and the assay was performed according to the manufacturer’s protocols. In brief, the PVDF membrane was coated with the IFN-γ capture antibody overnight, then washed. The antigens were plated at the specified concentrations in 100 μL/well. The PBMC were added at 300,000 cells per well in 100 μL CTL-Test™ Medium, and the plates were gently tapped on each side to ensure even distribution of the cells. After a 24 h incubation at 37 °C in a humidified CO2 incubator, during which the IFN-γ produced by the antigen-stimulated T cells was captured, the cells were discarded, IFN-γ detection antibody was added, and the spot forming units (SFU) were detected via enzyme-catalyzed substrate precipitation. The plates were air-dried prior to analysis. The plates were analyzed using an ImmunoSpot^®^ S6 Ultimate Reader from CTL (Catalog# S6UTM12). The numbers of SFU were established using the ImmunoSpot^®^ Software’s (from CTL) SmartCount™ and Autogate™ functions [28] that permit user-independent objective counting of SFUs [29]. Spot counts reported for the respective antigen-stimulated test conditions are means from triplicate wells, without the medium control subtracted.

### 2.5. Statistical Analysis of ImmunoSpot^®^ SFU Counts

As ImmunoSpot^®^ counts are normally distributed among replicate wells, the utilization of parametric statistics is suited for identifying positive responses [30]. Accordingly, the Student’s *t*-test was done comparing SFU counts in the triplicate antigen-containing wells *vs*. the SFU counts in the triplicate medium control wells. A *p*-value <0.05 was considered as the cut-off for a significant SFU increase.

## 3. Results

### 3.1. CEF and CERI Recall CD8+ T Cells, CPI CD4+ T Cells, CEFX Both

CD8+ T cells recognize 8-11 amino acid long peptides presented to them on HLA class I molecules [18]. As the peptide-binding groove of class I molecules is closed on both ends, it cannot accommodate longer peptides [31]. MHC class II molecules, in contrast, cannot efficiently bind and present such short peptides to CD4+ T cells [31]. As both the CEF and the CERI peptide pools contain peptides of 8-11 amino acid length, one would expect both to recall CD8+ T cells only. We performed cell depletion experiments to verify this assumption. As shown in Figure 1, CD8+ T cell-depleted PBMC fractions (PBMC-CD8) lost the CEF and CERI peptide pool-triggered recall response *vs*. the unseparated PBMC while CD4+ T cell depletion (PBMC-CD4), in contrast, had no such effect. These data suggest that the CEF- and CERI-triggered IFN-γ SFU are indeed produced by CD8+ T cells.

**Figure 1.**
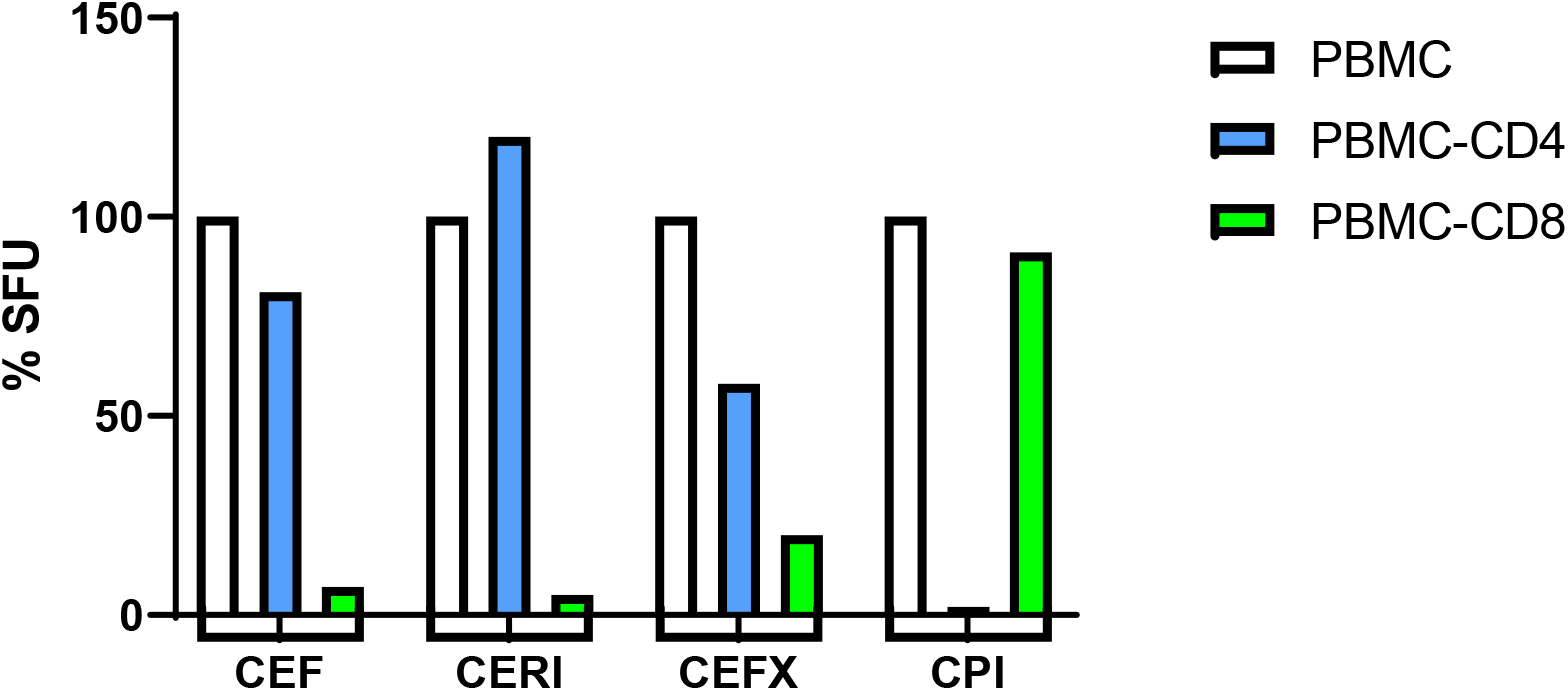
Identifying the CD4/CD8 lineage of T cells responding to antigens CEF, CERI, CEFX and CPI. The SFU counts in the unseparated PBMC was set as 100%, to which the SFU counts in the CD4 cell-depleted PBMC (PBMC-CD4) and the CD8 cell-depleted PBMC (PBMC-CD8) are compared. PBMC, PBMC-CD4, and PBMC-CD8 were all adjusted to 300,000 cells/well.

We also tested purified CD8+ T cells obtained from PBMC by negative selection. CEF and CERI activated IFN-γ SFU in these purified CD8+ T cell fractions without the need to add APC (data not shown), further supporting the notion that CD8+ T cells are responding to CEF and CERI, as CD8+ T cells are HLA class I-positive and they can serve as APCs to each other.

CD4+ T cells recognize protein antigens that are presented by specialized HLA class II-positive APC (primarily macrophages, B cells and dendritic cells) which internalize, process and present the antigen to the CD4+ T cells [32]. CPI consists of native viral proteins, and as such can be expected to be recognized by CD4+ T cells [26]. Cell separation experiments confirmed this notion: CD4+ T cell depletion from PBMC abrogated the CPI-triggered recall response, whereas CD8+ T cell depletion had no effect on it. When we tested purified CD4+ T cells (obtained from PBMC by negative selection) we found that the CPI-induced recall response could only be detected if APC were added to the purified CD4+ T cells (data not shown). CPI therefore recalls CD4+ T cells, and unlike CEF and CERI, requires antigen processing and presentation: therefore, CPI also tests the functionality of the APC compartment.

CEFX has been designed to be a universal positive control for the recall of CD4+ and CD8+ T cells alike, and accordingly consists of peptides 9 to 15 amino acids long that are suited for direct HLA class I and HLA class II molecule binding, without the need for additional antigen processing. When tested on the PBMC-CD4 and PBMC-CD8 cell fractions *vs*. the unseparated PBMC, an impairment was seen in both cell fractions, confirming a mixed recall of CD4+ and CD8+ T cells.

The data so far show that a single positive control might not suffice for the comprehensive assessment of PBMC fitness for T cell immune monitoring. CEF and CERI are candidates to test the functionality of antigen-specific CD8+ T cells, but do not serve to assess the functionality of the APC compartment. For testing the functionality of CD4+ T cells, and that of the antigen processing machinery, CPI is an ideal candidate. CEFX tests the functionality of CD4+ and CD8+ T cells, but owing to the 9-15 amino acid peptides that can be loaded directly into class I and class II molecules, it does not address the functionality of antigen processing.

### 3.2. CEF Fails as a Positive Control in 48% of Test Subjects

A positive control should ideally work for all test subjects. To verify whether this is the case for the CEF peptide pool, we tested all 210 healthy donors currently available in the ePBMC library. Standard 24 h IFN-γ ELISPOT assays were performed testing 300,000 PBMC per well. The raw data are shown in Supplemental Table 1, including medium control and CEF-triggered SFU counts and the class I HLA type, sex, age, and race of the test subjects. The CEF peptide pool-triggered SFU counts ranged between “too numerous to reliably count” (TNTC, > 500 SFU/well) and zero. From the perspective of a positive control, we argue that qualifying results should exceed 50 SFU/300,000 PBMC, because such response magnitudes are commonly seen with individual peptides or antigens [10]. Of the 210 donors tested, 100 (48%) fell in the < 50 SFU/300,000 PBMC category (Figure 2A). Forty nine of the 210 donors (23%) showed no CEF response (< 10 SFU/300,000 PBMC), and 51 subjects (24%) of all donors tested fell in the 10-49 SFU/300,000 PBMC category. The breakdown of CEF responses by race of the test subjects is shown in Figure 2B.

**Figure 2.**
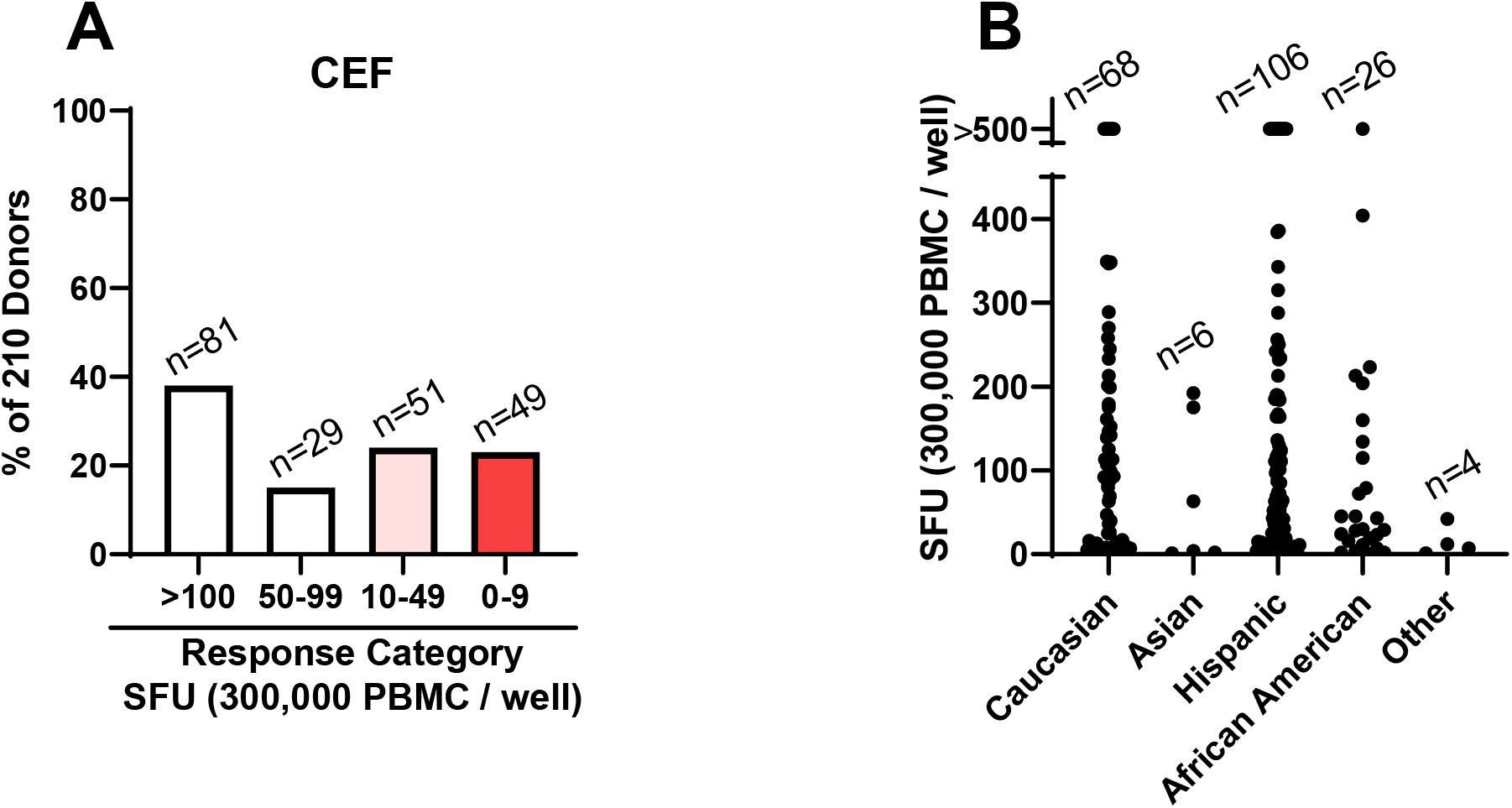
CEF peptide pool-triggered recall response in 210 healthy human donors. A standard 24 h IFN-γ ImmunoSpot^®^ assay was performed testing the CEF peptide pool induced SFU numbers. (A) Response magnitudes have been divided into the specified SFU categories. The percentage of subjects falling into each response category is shown on the Y axis, and the number of subjects in each category is specified above the bars. (B). The breakdown of CEF responses by race is shown with each dot representing the results obtained for one test subject.

### 3.3. CEF Non-Responder PBMC Respond to Anti-CD3 Stimulation

Non-responsiveness to a positive control could either mean that the PBMC are damaged/non-fit, or that the positive control itself is suboptimal. To distinguish between these two fundamentally different scenarios, we subjected samples of PBMC from CEF non-responder subjects to anti-CD3 stimulation. As listed in Table 1, with few exceptions, anti-CD3 stimulation triggered vigorous IFN-γ SFU formation in CEF non-responder PBMC. The only subject with an impaired PBMC response to anti-CD3 stimulation was ID 82. Reduced anti-CD3 triggered IFN-γ SFU formation using samples of ID 101, 118, and 131 also suggests that these PBMC might qualify as impaired too. However, the strong anti-CD3 responsiveness of all other CEF non-responder PBMC clearly establishes that it is not these PBMC’s impaired functionality, but instead the insufficient formulation of the CEF peptide pool that accounted for the CEF-negative result.

**Table 1.** CEF-negative PBMC can respond to other T cell stimuli. PBMC that showed no response to CEF (< 10 SFU/300,000 PBMC) were tested simultaneously for CERI, CEFX, CPI and anti-CD3-induced T cell activation in a standard 24 h IFN-γ ImmunoSpot^®^ assay at 300,000 PBMC/well, in triplicates each. Mean SFU counts are shown for all conditions.

| Subject | Media | CEF | CERI | CEFX | CPI | Anti-CD3 |
| --- | --- | --- | --- | --- | --- | --- |
| ID 30 | 1 | 8 | 117 | 96 | 212 | >500 |
| ID 51 | 2 | 7 | 138 | 41 | 268 | >500 |
| ID 68 | 2 | 2 | 413 | 265 | 270 | >500 |
| ID 69 | 0 | 0 | 2 | 4 | 6 | 436 |
| ID 71 | 2 | 4 | 51 | 160 | 352 | >500 |
| ID 82 | 1 | 1 | 15 | 27 | 4 | 129 |
| ID 87 | 2 | 1 | 48 | 17 | 158 | >500 |
| ID 98 | 2 | 3 | 252 | 118 | 250 | >500 |
| ID 101 | 1 | 6 | 72 | 29 | 29 | 233 |
| ID 103 | 1 | 1 | 112 | 180 | 194 | >500 |
| ID 118 | 0 | 2 | 32 | 18 | 212 | 243 |
| ID 123 | 2 | 4 | 180 | 58 | 366 | 348 |
| ID 128 | 1 | 7 | 121 | 53 | 183 | 459 |
| ID 131 | 1 | 9 | 47 | 23 | 37 | 260 |
| ID 133 | 3 | 2 | 159 | 89 | 316 | >500 |
| ID 138 | 1 | 5 | 32 | 123 | 461 | >500 |
| ID 144 | 0 | 7 | 55 | 127 | 166 | >500 |
| ID 147 | 1 | 5 | 77 | 156 | 500 | >500 |
| ID 150 | 0 | 8 | 83 | 410 | 216 | >500 |
| ID 160 | 1 | 2 | 12 | 43 | 27 | 495 |
| ID 168 | 3 | 6 | 33 | 136 | 439 | >500 |
| ID 169 | 2 | 8 | 102 | 70 | 293 | 381 |
| ID 191 | 1 | 0 | 70 | 75 | 154 | 403 |
| ID 196 | 1 | 5 | 32 | 20 | 154 | >500 |

### 3.4. CEF Non-/Low-Responder PBMC Respond to CERI, CEFX and CPI Stimulation

To further assess the fitness of CEF-non/low-responsive PBMC, we included CERI, CEFX, and CPI antigens into the functionality testing. Like anti-CD3, they too induced in part very high SFU numbers in the CEF non-responders (Table 1, and Figure 3A-C) and CEF low-responders as well (10-49 SFU, Figure 3D-F). These very strong antigen-specific CD8+ and CD4+ T cell recall responses in CEF-low/non-responding PBMC further establish that it is not these PBMC’s impaired functionality, but the insufficient formulation of the CEF peptide pool itself that accounts for the CEF-negative results.

**Figure 3.**
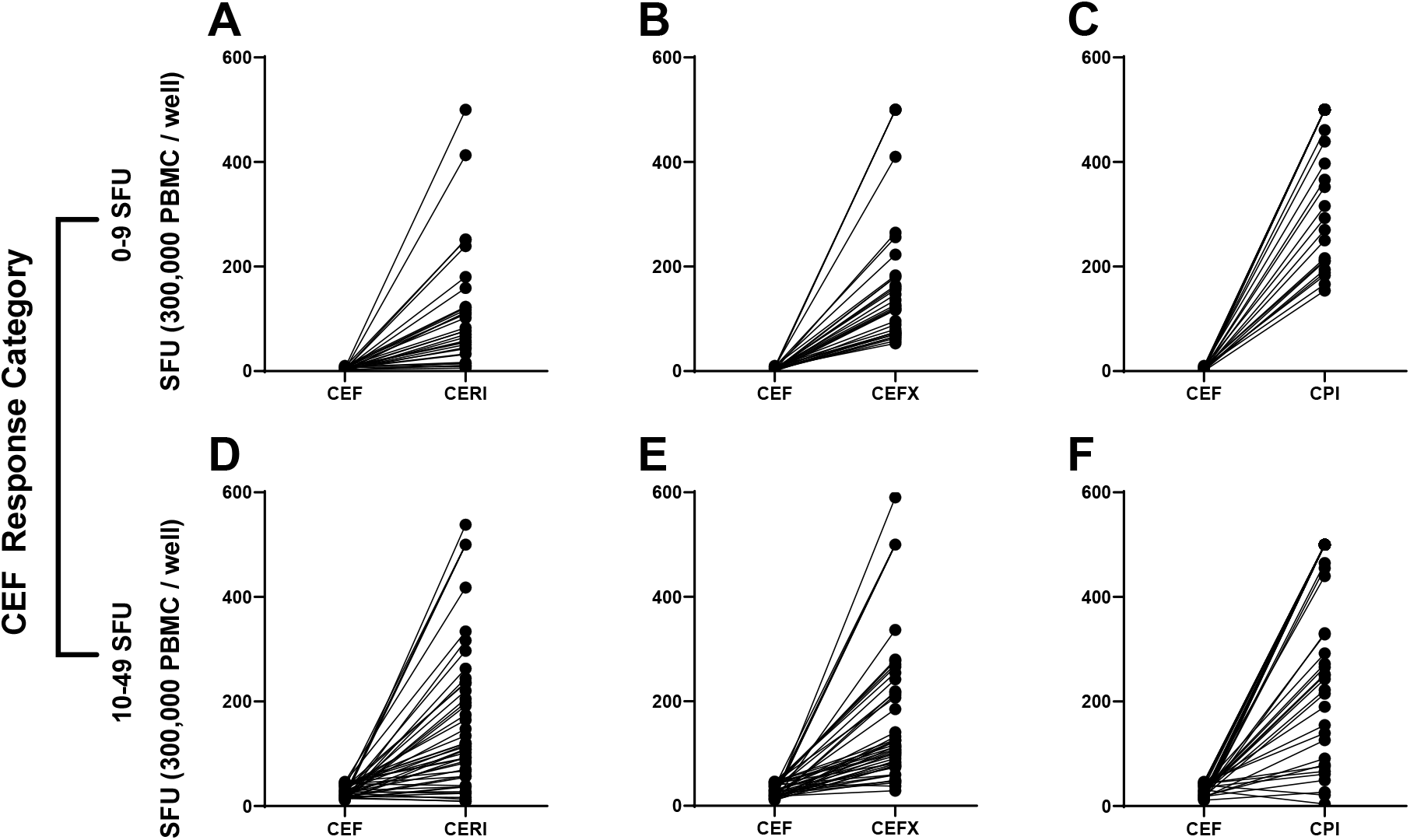
CERI-, CEFX-, and CPI-induced recall responses in CEF non-/low-responder PBMC. (A-C) Subjects whose PBMC fell with < 10 SFU/300,000 PBMC into the CEF non-responder category (n=49), or (D-E) with 10-49 SFU/300,000 PBMC into the CEF low-category (n=51). were tested in a standard 24 h IFN-γ ImmunoSpot^®^ assay for CERI (A&D), CEF-X (B&E) and CPI (C&F) recall at 300,000 PBMC/well. SFU counts of the same PBMC are connected with a line.

### 3.5. Low/Non-Responders to CERI, CEFX and CPI are Rare

Encouraged by the above findings, we set out to test all 210 subjects’ PBMC for recall responses to CERI, CEFX and CPI. As shown in Figure 4, only 4% of these PBMC were negative for CERI, <1% for CEFX, and 2% for CPI, respectively, compared to the 23% CEF-negatives (Figure 1). In the borderline/low category (10-49 CEF-induced SFU/300,000 PBMC), there were 13% for CERI, 10% for CEFX, and 5% for CPI, compared to the 24% of CEF-responders in this category. Genuine positive controls can be expected to induce a stronger recall response than individual antigens do, a threshold we empirically set at < 50 SFU/300,000 PBMC. For CEF, 52% (110/210) of the test subjects’ PBMC reached this threshold (see Figure 1). In contrast, this threshold was reached for CPI by 93% (195/210), for CEFX by 89% (187/210) and for CERI by 83% (174/210) of the test subjects’ PBMC.

**Figure 4.**
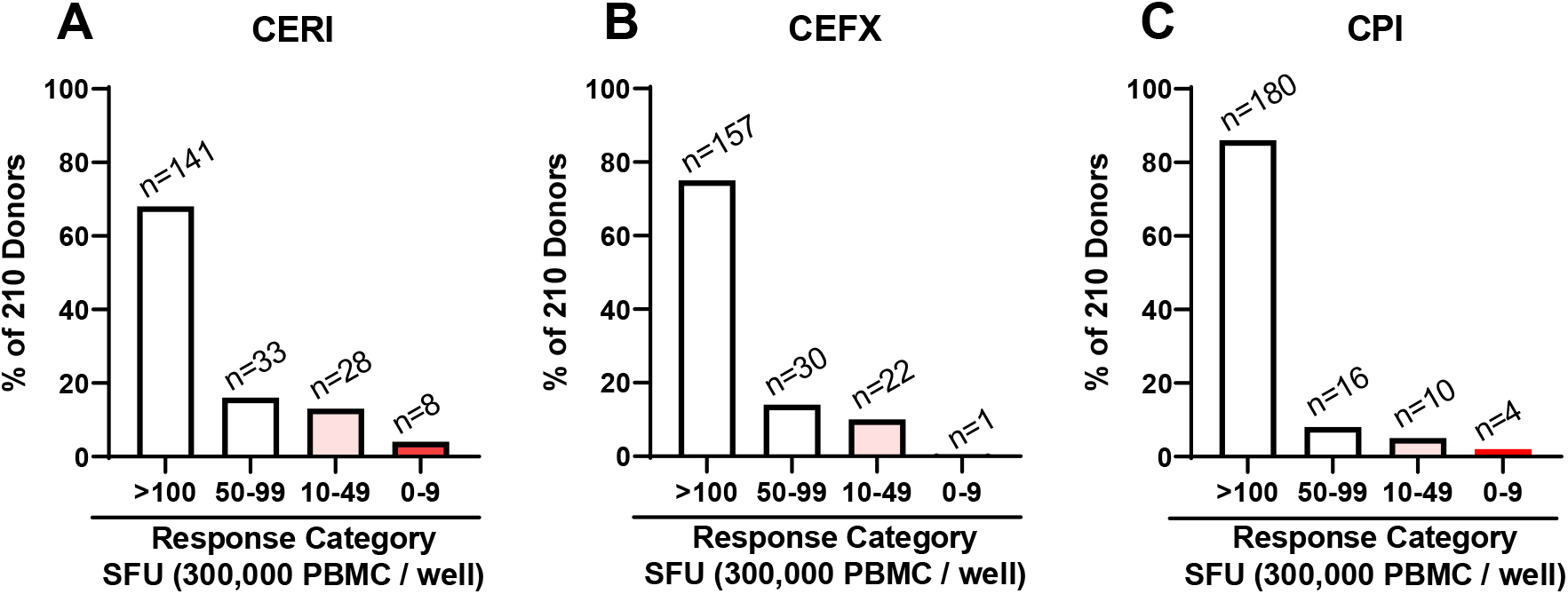
CERI, CEFX and CPI-triggered recall responses in 210 healthy human donors. A standard 24 h IFN-γ ImmunoSpot^®^ assay was performed testing the antigen-induced SFU numbers. Response magnitudes have been divided into the specified response categories. The percentage of subjects falling into each response category is shown on the Y axis, and the number of subjects in each category is specified above the bars.

## 4. Discussion

The data presented here suggest that the CEF peptide pool is a suboptimal positive control for testing the functionality of antigen-specific T cells in PBMC. CERI, CEFX and CPI by far outperform CEF in this respect. However, each of these positive controls tests a different T cell compartment. CERI, like CEF itself, tests the functionality of antigen-specific CD8+ T cells. As the short peptides contained in the CEF and CERI pools bind directly to HLA class I molecules expressed on all cell lineages present in PBMC, these peptide pools do not address the functionality of the antigen processing machinery. CPI, in contrast, consists of whole proteins that require antigen processing and presentation by professional APC present within PBMC, primarily macrophages, dendritic cells and B cells. CPI therefore tests for both CD4+ T cell and APC functionality. CEFX, consisting of peptides capable of direct binding to HLA class I and class II molecules, probes antigen-specific CD8+ and CD4+ T cell functionality; however, is unable to address functionality of the antigen processing compartment.

Viewing the data in Table 1, one might conclude that anti-CD3 is the ideal positive control for assessing PBMC fitness. Indeed, being a polyclonal CD4+ and CD8+ T cell stimulator, anti-CD3 activates *in vivo* differentiated type 1 (IFN-γ-producing) T cells in much higher frequency than individual antigens or pools of antigens can do. However, anti-CD3 antibodies result in unnatural T cell stimulation [33], that is fundamentally different from the serial triggering involved in regular T cell activation during which the T cell receptor (TCR) oscillates between low affinity binding to, and dissociation from, its ligand, the HLA-nominal peptide complex [34]. Therefore, in these authors’ eyes, in addition to anti-CD3, T cell immune monitoring is also in need of antigens that are suited to test physiologic T cell activation assessing the different requirements of CD4+ or CD8+ T cell detection in PBMC, including the APC’s functionality.

The data presented here show how difficult it is to come up with universal positive controls for T cell immune monitoring. CPI came closest to detect antigen-specific T cells in all donors tested, inducing > 50 SFU/300,000 PBMC in 93% of them. However, the response magnitude to CPI (and to all of the other positive control antigens tested) showed a wide span of inter-individual variations. This outcome is expected, as not all individuals have developed T cell immunity to all antigens contained in the respective positive control (in the case of CPI, CD4+ T cell responses to proteins from CMV, Parainfluenza, and Influenza viruses), and if they did, the frequencies of T cells targeting each of these viruses will differ among individuals dependent upon their immune status *vs*. the respective virus. Therefore, inclusion of antigens from additional viruses into the CPI panel might be required to elicit larger recall responses in the remainder (7%) of CPI low/non-responders.

When CPI reactivity is tested, the viral proteins are processed and presented by the autologous APC present in the PBMC. Therefore, the correct and complete selection of epitopes displayed to CD4+ T cells on APC is not a limiting factor; it occurs naturally. Thus, it can be assumed that the *ex vivo* recall with CPI assesses the entire *in vivo* primed CPI-specific CD4+ T cell repertoire. However, when peptides are used for recall, as in the case of CEF, CERI and CEFX, the peptides used for recall are not likely to address the entire virus-specific T cell repertoire.

The underlying assumption for creating the CEF pool was that there is immune dominant recognition of a few epitopes from CMV, EBV and influenza viruses. For example, it was assumed that a single peptide of CMV, pp65_495-503_, is immune dominant in all HLA-A*02:01-positive subjects and therefore this peptide would suffice to detect CMV-specific CD8+ T cells in all CMV-infected, HLA-A*02:01-positive humans [8]. In a recent study, complete epitope mapping was performed for the CMV pp65 protein on four CMV-positive, HLA-A*02:01-positive test subjects [35]. Peptide pp65_495-503_ was indeed immune dominant in one of these donors, but it was cryptic (it induced a borderline/low recall response) in another HLA-A*02:01-positive, CMV-positive donor, who in turn exhibited dominant CD8+ T cell recognition of two alternative pp65-derived epitopes. In yet two other HLA-A*02:01-positive, CMV-positive donors, pp65_495-503_ was subdominant, recalling low frequency SFU compared to other dominant epitopes of the pp65 antigen. Therefore, upon closer examination, the immune dominance of pp65_495-503_ does not hold up for HLA-A02:01-positive subjects. Rather pp65_495-503_ is just one of several potential CD8+ T cell epitopes to which CD8+ T cells respond in an apparently aleatory (dice-like) manner [19].

The above conclusion was also supported by a study [10] in which the individual CEF peptides, including pp65_495-503_, were systematically tested on high-resolution HLA-typed healthy test subjects. It was found that of 241 expected positive recall responses, in only 36 (15%) instances did the expected individual CEF peptides indeed recall strongly positive (dominant) CD8+ T cell responses, in 41 (17%) instances they induced low frequency CD8+ T cells (subdominant), and in 68% of the test cases these expectedly immune dominant epitopes recalled a borderline or negative (cryptic) CD8+ T cell response. Similar results were obtained for the CMV pp65_495-503_ peptide in HLA-A*02:01-positive, CMV-infected donors in the aforementioned study[10]. This observation was confirmed in a follow-up study that involved 52 HLA-A*02:01-positive, CMV-infected subjects: even though all these subjects have developed strong T cell immunity to CMV, 8% of them did not respond to the CMV pp65_495-503_ peptide at all, and 27% displayed subdominant/cryptic response to the pp65_495-503_ peptide[23].

Immune dominance of single epitopes might therefore be the exception even in HLA-allele matched humans, and aleatory recognition of multiple epitopes the rule [19], suggesting that positive controls that rely on a few select peptides are prone to underestimate, or outright miss, the virus-specific memory T cells they were designed to detect. The 32 peptide-containing CEF pool contains only five CMV peptides attempting to detect CMV-specific CD8+ T cells across the human population The shortcoming of the CEF peptide pool is therefore linked to the absence of immune dominance. Thus, due to the tremendous HLA class I allele diversity in the human population, these 32 peptides of CMV, EBV and Flu virus are insufficient to reliably detect CD8+ T cells specific for these viruses. Relying on 176 known epitopes of a larger diversity of viruses, and 124 peptides of predicted epitopes, the CEFX and CERI peptide pools come much closer, but still do not completely meet the goal of all-encompassing positive controls. The long-term solution will be to further increase the number of antigens and peptides in positive controls. The short-term solution is to use all three highly improved positive control antigens presently available: CEFX, CERI and CPI, in addition to anti-CD3. Due to the response magnitudes elicited by these, it is not necessary to test each in replicates, and therefore all three positive control antigens can be tested with the same numbers of PBMC as presently done when CEF is tested in triplicate. Testing of all three positive controls is not only advisable because they are complementary in covering a wider spectrum of recall antigens, to each of which test subjects are prone to have various levels of T cell immunity, but also because in this way the functionality of CD4+, CD8+ T cells, and of the APC are separately addressed, which makes a comprehensive assessment of PBMC fitness possible.

## Acknowledgments

We would like to thank Ruliang Li for expert technical assistance, Kyle Schulz for advice in regulatory matters, and Diana Roen and Greg Kirchenbaum for editorial assistance. This study was conducted in the R&D department of CTL, and was fully funded from the research budget of CTL.

## Author Contributions

P.V.L. and A.L conceived and designed the experiments; P.A.R. designed the CERI peptide library. A.L. performed all experiments with assistance of T.Z. and M.S. The manuscript was written by P.V.L and A.L. This publication serves as part of A.L.’s doctoral thesis to be submitted to the Universidad Complutense de Madrid, Madrid, Spain.

## Sample Availability

All samples used are commercially available products.

## References

1. Schumacher, T.N.; Schreiber, R.D. Neoantigens in cancer immunotherapy. Science 2015, 348, 69–74, doi:10.1126/science.aaa4971

2. Sette, A.; Crotty, S. Pre-existing immunity to SARS-CoV-2: the knowns and unknowns. Nature Reviews Immunology 2020, 20, 457–458, doi:10.1038/s41577-020-0389-z.

3. Tough, D.F.; Sprent, J. Life span of naive and memory T cells. Stem Cells 1995, 13, 242–249, doi:10.1002/stem.5530130305.

4. Kuerten, S.; Batoulis, H.; Recks, M.S.; Karacsony, E.; Zhang, W.; Subbramanian, R.A.; Lehmann, P.V. Resting of cryopreserved PBMC does not generally benefit the performance of antigen-specific T cell ELISPOT assays. Cells 2012, 1, 409–427. doi:10.3390/cells1030409.

5. Prabhakar, U.; Kelley, M. Validation of cell-based assays in the GLP setting: a practical guide; Wiley; Chichester: John Wiley [distributor]: Hoboken, N.J., 2008.

6. Kreher, C.R.; Dittrich, M.T.; Guerkov, R.; Boehm, B.O.; Tary-Lehmann, M. CD4+ and CD8+ cells in cryopreserved human PBMC maintain full functionality in cytokine ELISPOT assays. Journal of Immunological Methods 2003, 278, 79–93, doi: https://doi.org/10.1016/S0022-1759(03)00226-6.

7. Wunsch, M.; Caspell, R.; Kuerten, S.; Lehmann, P.V.; Sundararaman, S. Serial measurements of apoptotic cell numbers provide better acceptance criterion for PBMC quality than a single measurement prior to the T cell assay. Cells 2015, 4, 40–55, doi:10.3390/cells4010040.

8. Currier, J.R.; Kuta, E.G.; Turk, E.; Earhart, L.B.; Loomis-Price, L.; Janetzki, S.; Ferrari, G.; Birx, D.L.; Cox, J.H. A panel of MHC class I restricted viral peptides for use as a quality control for vaccine trial ELISPOT assays. Journal of Immunological Methods 2002, 260, 157–172, doi:10.1016/s0022-1759(01)00535-x.

9. Geiger, R.; Duhen, T.; Lanzavecchia, A.; Sallusto, F. Human naive and memory CD4+ T cell repertoires specific for naturally processed antigens analyzed using libraries of amplified T cells. The Journal of Experimental Medicine 2009, 206, 1525–1534, doi:10.1084/jem.20090504.

10. Moldovan, I.; Targoni, O.; Zhang, W.; Sundararaman, S.; Lehmann, P.V. How frequently are predicted peptides actually recognized by CD8 cells? Cancer Immunology, Immunotherapy, 2016, 65, 847–855, doi:10.1007/s00262-016-1840-7.

11. Chen, W.H.; Strych, U.; Hotez, P.J.; Bottazzi, M.E. The SARS-CoV-2 vaccine pipeline: an overview. Current Tropical Medicine Reports 2020, 10.1007/s40475-020-00201-6, 1–4, doi:10.1007/s40475-020-00201-6.

12. Lehmann, P.V.; Zhang, W. Unique strengths of ELISPOT for T cell diagnostics. Methods in Molecular Biology (Clifton, N.J.) 2012, 792, 3–23, doi:10.1007/978-1-61779-325-7_1.

13. Hesse, M.D.; Karulin, A.Y.; Boehm, B.O.; Lehmann, P.V.; Tary-Lehmann, M. A T cell clone’s avidity is a function of its activation state. The Journal of Immunology 2001, 167, 1353–1361, doi:10.4049/jimmunol.167.3.1353.

14. Hanson, J.; Roen, D.R.; Lehmann, P.V. Four color ImmunoSpot^®^ assays for identification of effector T cell lineages. Methods in Molecular Biology (Clifton, N.J.) 2018, 1808, 51–62, doi:10.1007/978-1-4939-8567-8_5.

15. Ramachandran, H.; Laux, J.; Moldovan, I.; Caspell, R.; Lehmann, P.V.; Subbramanian, R.A. Optimal thawing of cryopreserved peripheral blood mononuclear cells for use in high-throughput human immune monitoring studies. Cells 2012, 1, 313–324, doi:10.3390/cells1030313.

16. Neefjes, J.; Jongsma, M.L.; Paul, P.; Bakke, O. Towards a systems understanding of MHC class I and MHC class II antigen presentation. Nature Reviews, Immunology 2011, 11, 823–836, doi:10.1038/nri3084.

17. Pamer, E.; Cresswell, P. Mechanisms of MHC class I-restricted antigen processing. Annual Review of Immunology 1998, 16, 323–358, doi:10.1146/annurev.immunol.16.1.323.

18. Reche, P.A.; Reinherz, E.L. Sequence variability analysis of human class I and class II MHC molecules: functional and structural correlates of amino acid polymorphisms. Journal of Molecular Biology 2003, 331, 623–641, doi:10.1016/s0022-2836(03)00750-2.

19. Lehmann, P.V.; Lehmann, A.A. Aleatory epitope recognition prevails in human T cell responses? 2020, 40, 225–235, doi:10.1615/CritRevImmunol.2020034838.

20. Reynisson, B.; Alvarez, B.; Paul, S.; Peters, B.; Nielsen, M. NetMHCpan-4.1 and NetMHCIIpan-4.0: improved predictions of MHC antigen presentation by concurrent motif deconvolution and integration of MS MHC eluted ligand data. Nucleic Acids Research 2020, 48, W449–w454, doi:10.1093/nar/gkaa379.

21. Mei, S.; Li, F.; Leier, A.; Marquez-Lago, T.T.; Giam, K.; Croft, N.P.; Akutsu, T.; Smith, A.I.; Li, J.; Rossjohn, J., et al. A comprehensive review and performance evaluation of bioinformatics tools for HLA class I peptide-binding prediction. Briefings in Bioinformatics 2020, 21, 1119–1135, doi:10.1093/bib/bbz051.

22. Brennick, C.A.; George, M.M.; Srivastava, P.K.; Karandikar, S.H. Prediction of cancer neoepitopes needs new rules. Seminars in Immunology 2020, 47, 101387, doi:10.1016/j.smim.2020.101387.

23. Lehmann, A.A.; Zhang, T.; Reche, P.A.; Lehmann, P.V. Discordance between the predicted vs. the actually recognized CD8+ T cell epitopes of HCMV pp65 antigen and aleatory epitope dominance. bioRxiv 2020, 10.1101/2020.11.06.371633, 2020.2011.2006.371633, doi:10.1101/2020.11.06.371633.

24. Grifoni, A.; Weiskopf, D.; Ramirez, S.I.; Mateus, J.; Dan, J.M.; Moderbacher, C.R.; Rawlings, S.A.; Sutherland, A.; Premkumar, L.; Jadi, R.S., et al. Targets of T cell responses to SARS-CoV-2 coronavirus in humans with COVID-19 disease and unexposed Individuals. Cell 2020, 181, 1489–1501.e1415, doi:10.1016/j.cell.2020.05.015.

25. Program, N.A.R. CEF control peptide pool NIH, Ed. aidreagent.org, 2018, https://www.aidsreagent.org/reagentdetail.cfm?t=peptides&id=291

26. Schiller, A.; Zhang, T.; Li, R.; Duechting, A.; Sundararaman, S.; Przybyla, A.; Kuerten, S.; Lehmann, P.V. A positive control for detection of functional CD4 T cells in PBMC: the CPI pool. Cells 2017, 6, doi:10.3390/cells6040047.

27. Molero-Abraham, M.; Lafuente, E.M.; Flower, D.R.; Reche, P.A. Selection of conserved epitopes from hepatitis C virus for pan-populational stimulation of T-cell responses. Clinical & Developmental Immunology 2013, 2013, 601943, doi:10.1155/2013/601943.

28. Karulin, A.Y.; Karacsony, K.; Zhang, W.; Targoni, O.S.; Moldovan, I.; Dittrich, M.; Sundararaman, S.; Lehmann, P.V. ELISPOTs produced by CD8 and CD4 cells follow log normal size distribution permitting objective counting. Cells 2015, 4, 56–70, doi:10.3390/cells4010056.

29. Zhang, W.; Lehmann, P.V. Objective, user-independent ELISPOT data analysis based on scientifically validated principles. Methods in Molecular Biology (Clifton, N.J.) 2012, 792, 155–171, doi:10.1007/978-1-61779-325-7_13.

30. Karulin, A.Y.; Caspell, R.; Dittrich, M.; Lehmann, P.V. Normal distribution of CD8+ T-cell-derived ELISPOT counts within replicates justifies the reliance on parametric statistics for identifying positive responses. Cells 2015, 4, 96–111, doi:10.3390/cells4010096.

31. Bjorkman, P.J.; Saper, M.A.; Samraoui, B.; Bennett, W.S.; Strominger, J.L.; Wiley, D.C. Structure of the human class I histocompatibility antigen, HLA-A2. Nature 1987, 329, 506–512, doi:10.1038/329506a0.

32. Rock, K.L.; Reits, E.; Neefjes, J. Present yourself! By MHC class I and MHC class II molecules. Trends in Immunology 2016, 37, 724–737, doi:10.1016/j.it.2016.08.010.

33. Norman, D.J. Mechanisms of action and overview of OKT3. Therapeutic Drug Monitoring 1995, 17, 615–620, doi:10.1097/00007691-199512000-00012.

34. Huppa, J.B.; Axmann, M.; Mörtelmaier, M.A.; Lillemeier, B.F.; Newell, E.W.; Brameshuber, M.; Klein, L.O.; Schütz, G.J.; Davis, M.M. TCR–peptide–MHC interactions in situ show accelerated kinetics and increased affinity. Nature 2010, 463, 963–967, doi:10.1038/nature08746.

35. Lehmann, P.V.; Suwansaard, M.; Zhang, T.; Roen, D.R.; Kirchenbaum, G.A.; Karulin, A.Y.; Lehmann, A.; Reche, P.A. Comprehensive evaluation of the expressed CD8+ T cell epitope space using high-throughput epitope mapping. Frontiers in Immunology 2019, 10, 655, doi:10.3389/fimmu.2019.00655.

